# Positive Selection on Rare Variants Underlying the Cold Adaptation of Wild Boar

**DOI:** 10.1101/2024.04.07.588424

**Authors:** Jianhai Chen, Ivan Jakovlić, Mikhail Sablin, Shengqian Xia, Zhixiang Xu, Yapin Guo, Renzuo Kuang, Jie Zhong, Yangying Jia, Thuy Nhien Tran Thi, Hao Yang, Hong Ma, Nikica Šprem, Jianlin Han, Di Liu, Yunxia Zhao, Shuhong Zhao

## Abstract

The wide geographical distribution of Eurasian wild boar (*Sus scrofa*) offers a natural experiment to study the thermoregulation. Here, we conducted whole-genome resequencing and chromatin profiling experiments on the local populations from cold regions (northern and northeastern Asia) and warm regions (southeastern Asia and southern China). Using genome-wide scans of four methods, we detected candidate genes underlying cold-adaptation with significant enrichment of pathways related to thermogenesis, fat cell development, and adipose tissue regulation. We also found two enhancer variants under positive selection, an intronic variant of *IGF1R* (rs341219502) and an exonic variant of *BRD4* (rs327139795), which showed the highest differentiation between cold and warm region populations of wild boar and domestic pigs. Moreover, these rare variants were absent in outgroup species and warm-region wild boar but nearly fixed in cold-region populations, suggesting their *de novo* origins in cold-region populations. The experiments of CUT&Tag chromatin profiling showed that rs341219502 of *IGF1R* is associated with the gain of three novel transcription factors involving regulatory changes in enhancer function, while rs327139795 of *BRD4* could result in the loss of a phosphorylation site due to amino acid alteration. We also found three genes (*SLCO1C1, PDE3A*, *and TTC28*) with selection signals in both wild boar and native human populations from Siberia, which suggests convergent molecular adaptation in mammals. Our study shows the adaptive evolution of genomic molecules underlying the remarkable environmental flexibility of wild boar.

## Introduction

One of the most fundamental questions in evolution is to understand how populations could adapt new environment (1, 2). The peripheral or marginal population, which inhabits the boundary of a species’ distribution area, is a valuable model for us to understand this question. Peripheral populations often migrate from their ancestral territories to adapt to new niches through population expansion. These populations face harsher and sometimes insurmountable environmental conditions at boundary zones, such as extreme low temperatures and a scarcity of food resources, which impede their further territorial expansion. The adaptation in challenging survival conditions make them ideal natural experiments for investigating the genetic bases of novel adaptive strategies in the face of environmental constraints.

Advancements in sequencing technology and population genetics are increasingly enabling the identification of functional genetic variants under natural selection for organisms to adapt new environments (3-8). Strong selective forces in derived populations would leave distinguished genomic signals segregating from source populations (2, 9). To discern selective signals in peripheral populations, studies often explore the patterns of multiple genomic parameters, including allele frequencies (2, 10-12), nucleotide diversity (13-15), and haplotype segregation (16, 17). For example, human populations have adapted to diverse environments, ranging from the tropical African regions to the cold peripheral regions of Siberia. Genome-wide scan based on patterns of haplotype and allele frequency in Siberian populations reveals selective signals of genes underlying cold adaptation (3). By exploring variants of these candidate genes, a globally low-frequency nonsynonymous variant in *CPT1A* has been identified as the most likely causative mutation, due to its high allele frequency in local Siberian populations (18).

In this study, we focus on the wild boar, *Sus scrofa*, a species with remarkable adaptive capabilities and wide distribution across diverse climatic regions in Eurasia. Their ecological niches encompass the humid tropics of Southeast Asia, extend through temperate zones, and reach the extreme area of the Qinghai-Tibet plateau as well as subarctic Siberia (19, 20). Phylogenetic analyses suggest that the Eurasian wild boar diverged from a clade of closely related *Sus* species at the onset of the Pliocene epoch (5.3-3.5 million years ago) within the tropical Asia (21). Subsequently, between 1 and 2 million years ago, the species expanded beyond tropical Asia across a substantial range of Eurasia, establishing various geographically distinct populations (19). Biogeographic studies have revealed a migratory trajectory of the wild boar from southern to northern Asia (22, 23). This long-range migration of wild boar suggests that populations from tropical and Siberian regions are source and peripheral populations, respectively. Thus, wild boar could serve as a natural animal model to investigate genetic bases subject to positive selection in response to novel climatic challenges under cold environment.

Following their migration from tropical Asia, wild boar have established their northernmost natural habitat in Siberia, reaching as far as 61°N (20). Given the species’ tropical origins, we hypothesize that the genomes of cold-region populations could exhibit signals of adaptation to colder climates. However, this hypothesis has not yet been empirically tested (20). Here, we conducted the whole-genome sequencing of wild boar populations from both tropical Asian and Siberian regions. We identified candidate genes and related biological pathways underlying cold adaptation of wild boar. Based on these selective genes, we detected the leading variants with the highest cold-warm differentiation and site-level signal of selective sweep among all regulatory and missense variants. We further explored functional implications of the leading variants with experiments of the Cleavage Under Targets and Tagmentation (CUT&Tag). Our study provides insights into the positively selected genes and rare variants potentially underlying the bioclimatic adaption of cold-region wild boar.

## Results

### The phylogenetic origin of wild boar from cold and warm regions

We obtained a total of 821 Gb whole-genome data from 11 new samples: six wild boar from Siberia and five from Southeast Asia (Figure 1a). After mapping to the pig genome reference (Sscrofa v11.1), the average sequencing depth was estimated to be 28.32x. We compiled two datasets for different analytical purposes, featuring varied sample sizes: a core dataset comprising 63 genomes, and an extended dataset encompassing 488 genomes (Supplementary Tables 1 and 2). Using a distance matrix based on identity by state (IBS) and the neighbor-joining method, we initially evaluated the phylogenetic relationships for all 488 samples (Figure 1b). We revealed that eastern Siberian samples (E, F, and G in Figure 1a) clustered within the clade of wild boar populations from northern China, northeastern China, and northeastern Asia (South Korea and Japan) (D), while the western Siberian samples (H in Figure 1a) clustered with the European wild boar. The southeastern Asian wild boar clustered close to the wild boar and indigenous breeds from southern China.

**Figure 1.**
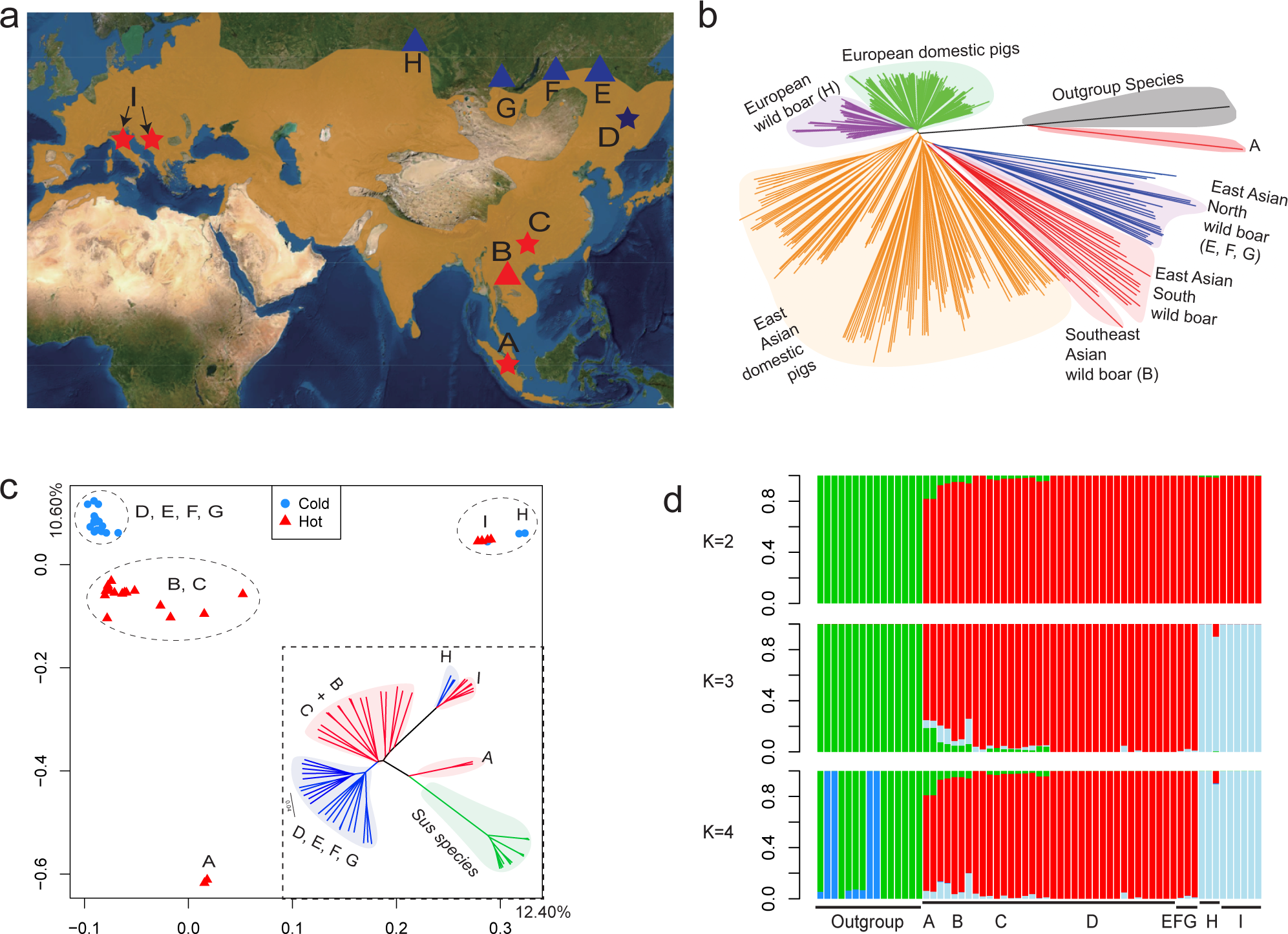
The population distribution, genomic relationships, and historical demography of wild boar populations. (a) The wild boar population distribution (map retrieved from the IUCN red list: https://www.iucnredlist.org/species/41775/44141833). The red and blue colors indicate samples from warm and cold regions, respectively. Triangles and stars indicate populations with newly sequenced data and publicly available data, respectively. The population locations are Sumatra (A), Vietnam (B), South China (C), Northeast China and Korea (D), Zabaykalsky Krai (E), Buryat (F), Tyva (G), Novosibirsk (H), and Italy and Greece (I). (b) The phylogenetic tree for the extended dataset of 448 samples is based on IBS distances among samples. Background shadows highlight major clades, and brackets indicate newly sequenced samples. (c) The principal component analysis (PCA) and phylogenetic tree based on autosomal SNPs. In the phylogenetic tree, all major clade divisions were supported by 1000 bootstrap replicates (100%). (d) The admixture analysis (K=2, 3, 4) for population ancestry. The green color indicates the outgroup *Sus* species (*S. verrucosus, S. celebensis, S. cebifrons,* and *S. barbatus*) from the islands of Southeast Asia. The red symbols show the Southeast and East Asian population ancestry in populations from A to G. The light blue shows the European ancestry in H and I.

For the core dataset, we conducted the principal component analysis (PCA) and IBS phylogenetic inference to validate population relationships (Figure 1c). These analyses revealed similar population phylogeny with the relationship obtained with the extended dataset (Figure 1b). Specifically, our newly sequenced tropical population (B) clustered together with the temperate wild boar from southern China (C in Figure 1). The newly sequenced Siberian population was divided into two clusters (E, F, and G vs. H, Figure 1c), consistent with the pattern produced by the extended dataset (Figure 1b). These results revealed higher divergence between the populations from western (H) and eastern Siberia (E, F, and G), but close relationship between tropical (B, Vietnam) and temperate Asian wild boar populations (southern China, C). These results also indicated a substantial genomic differentiation between peripheral populations of cold regions and source populations in tropical regions.

Gene flow between the western and eastern Siberian populations was examined utilizing TreeMix (v1.12) (24) (Supplementary Figure 1b-d). This analysis identified the optimal number of migration events (m) as three, which explained over 99.8% of the variance in genetic relatedness among the populations (Supplementary Figure 1b-c). Notably, the inferred migration events suggested a predominant westward gene flow from eastern to western Siberian populations (Supplementary Figure 1d). Further analysis on population structure was conducted using ADMIXTURE v1.3 (25), with a cross-validation approach pinpointing the optimal number of distinct ancestries at K=4 (Supplementary Figure 1a). An investigation into the ancestral composition, considering ancestry counts from two to four, distinguished two primary clades correlating with the established ancestries of Eurasian wild boar and domestic pigs, namely Asian and European lineages (Figure 1d). This division was corroborated by Principal Component Analysis (PCA) and phylogenetic assessments, elucidating the major population relationships (Figures 1b and 1c). Intriguingly, the population in western Siberia exhibited a 9.64% composition of Asian ancestry when analyzed at K=3 (Figure 1d), suggesting a restricted gene flow from Asia towards the western Siberian demographic.

### The genes under selective sweep and their functional enrichment in thermogenic and adipose-related pathways

Extensive research has suggested that 1% - 15% of genes in the mammalian genome may undergo positive Darwinian selection (26-28). For instance, in humans, approximately 10% of genes are believed to be under positive selection, though the concordance rates among different statistical tests typically range from 8% to 27% (27, 28). In our study, employing four complementary analytical approaches (Materials and Methods), we identified candidate genes showing evidence of selective pressure in cold-region wild boar, including those native to Siberia, northern and northeastern China, and northeastern Asia (Supplementary Table 2-7). Our comprehensive analysis revealed that 1.54% of genes (313 out of 20,306, Ensembl v105) were consistently identified across at least three distinct methods as being under selection, indicating a conservative estimation in this study. Additionally, 0.45% of genes (92 out of 20,306) were corroborated by all four analytical methods (Supplementary Table 7).

We analyzed the functional enrichment for genes supported by at least three methods (Figure 2a, Supplementary Table 8). We revealed four pathways potentially related to cold resistance: the ‘thermogenesis’ pathway (*ADCY9*, *NDUFB6*, *PPARG*, PRKACB, *SMARCC1*, *TSC2*), the ‘regulation of cold-induced thermogenesis’ pathway (*ACADL*, *IGF1R*, *JAK2*, *NOVA1*, *NOVA2*), the ‘positive regulation of adipose tissue development’ pathway (*PPARG*, *NCOA2*, *SIRT1*), and the ‘fat cell differentiation’ pathway (*PPARG*, *TGFB1*, *SIRT1*, *WWTR1*) (*p* < 0.05, Figure 2a). These findings indicate that specific pathways—especially those governing thermogenesis and fat cell differentiation—may play a critical role in enabling cold-region wild boars to adapt to their harsh environmental conditions.

**Figure 2.**
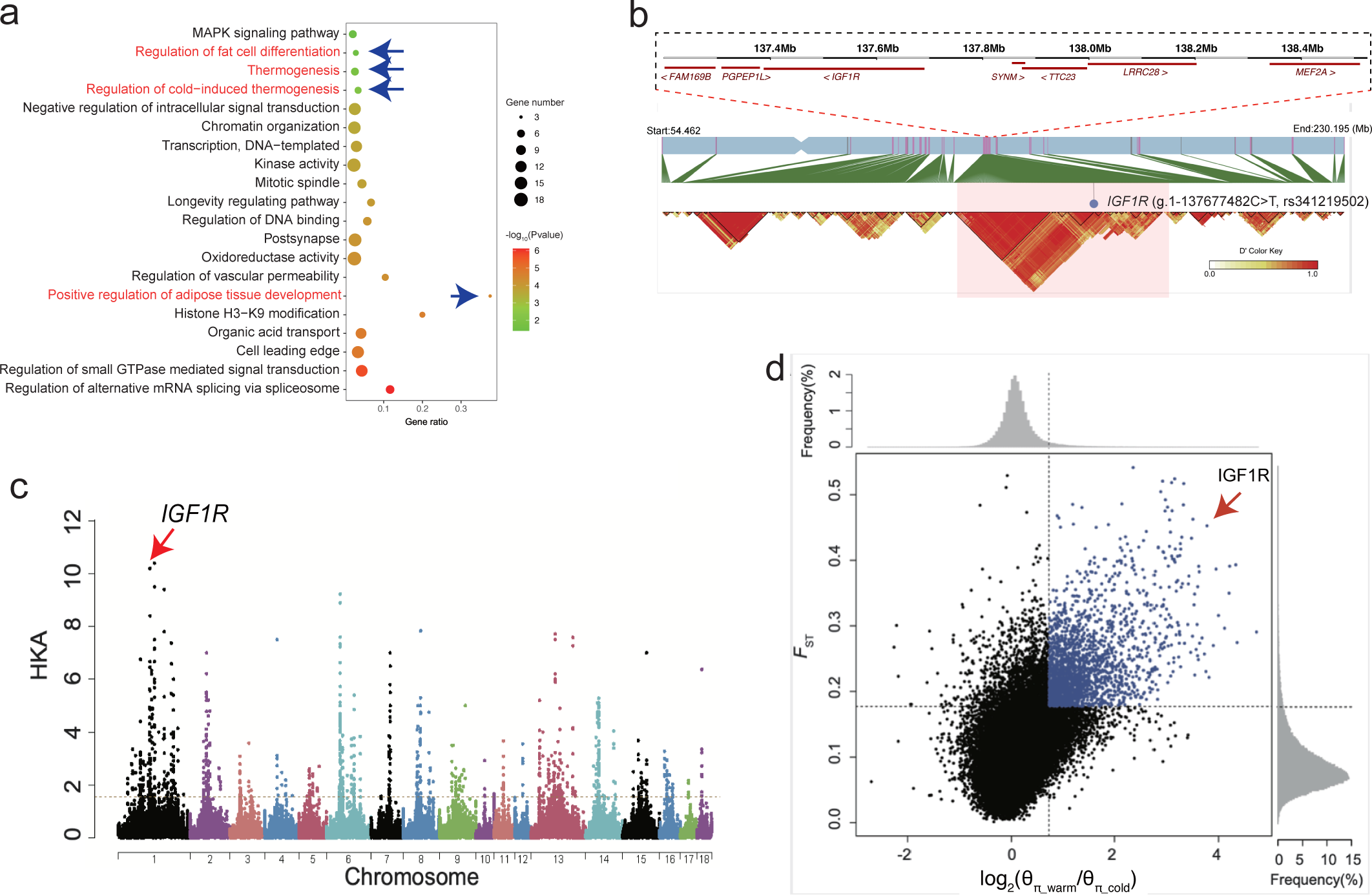
The functional enrichment and the candidate gene *IGF1R* underlying cold adaptation in the cold-region wild boar. (a) The enriched pathways were analyzed using the Metascape database (16). Only the significantly enriched pathways (*p*<0.05) are listed (Supplementary table 8). (b) Haplotype blocks for the cold and warm region samples based on variants with allele frequencies higher in cold than in warm regions (*ΔdAF*_cold-warm_ > 0.5). (c) The plot of HKA-like test. The vertical axis shows the decreased numbers of polymorphic sites relative to the counts of interspecies divergent sites. (d) The distribution of the Fixation index (*F_st_*) and nucleotide diversity ratio [log2(θ_π_warm_/θ_π_cold_)].

By analyzing genetic variants whose allele frequencies were higher in wild boar from cold region (Siberia, Korea, northern and northeastern China) than in those from warmer regions (temperate and tropical Asia), we uncovered a long haplotype block with strong linkage disequilibrium (LD, *ΔdAF_cold-warm_* > 0.5, D’ > 0.9) among variants of seven genes (*FAM169B*, *PGPEP1L*, *IGF1R*, *SYNM*, *TTC23*, *LRRC28*, and *MEF2A*). Surprisingly, this region is the longest gene cluster we identified with selective signals, spanned 1.3 Mb on chromosome 1 (137.2 Mb-138.5 Mb, Figure 2b). Among these linked genes, *IGF1R*, known as the Insulin-like Growth Factor 1 Receptor, received support from all four methods (Supplementary Table 7), suggesting the reliability of positive selection on this gene. Using the HKA-like test (29), we identified *IGF1R* as the most distinguished gene with the highest reduction of polymorphism relative to divergence, supporting a significant deviation from neutral evolution (Figure 2c). At population level, *IGF1R*, *ALDH1A2*, and *PGPEP1L* showed the highest inter-population divergences between cold and warm region wild boar among all protein-coding genes (*F_st_* = 0.65, Figure 2d and Supplementary Table 3). Notably, among these three genes, *IGF1R* demonstrated the highest reduction in nucleotide diversity for cold region wild boar compared to warm region ones (Figure 2d), suggesting a strong positive selection on linked variants beneficial to cold adaptation. These results of natural selection on *IGF1R* are consistent with the extensive *in vivo* studies on mice, which have revealed the role of *IGF1R* in thermoregulation, particularly in reducing core body temperature in response to cold stress and calorie restriction (30, 31).

### The potential convergent evolution for Siberian mammals

The endothermic mammalian species inhabiting cold regions like Siberia may have shared genes under convergent selection for cold resistance. To test this hypothesis, we retrieved genes under positive selection for cold adaptation in Siberian human populations, which were inferred with iHS and XP-EHH methods (top 1% windows) in a previous study (3) (Supplementary Table 9). We focused on shared genes with rigorous support from four methods (Supplementary Table 7), and found three consensus genes, *SLCO1C1, PDE3A*, *and TTC28*, which could be under convergent evolution. We confirmed the signals of positive selection for these genes based on nucleotide diversity comparison between cold- and warm-region populations and the HKA test (Supplementary Figure 2).

### The most differentiated intronic and exonic variants were detected in *IGF1R* and *BRD4*, respectively

Natural selection, driven by emerging selective forces, can result in the increased frequency or fixation of derived alleles, as well as the decreased frequency or loss of ancestral alleles within a peripheral population due to novel adaptation (18, 32). The causative variants within genes under positive selection are expected to show pronounced allele frequency divergence between peripheral and source populations (18). This principle has also been applied extensively in medical genetics to find disease variants (33-35). By analyzing polymorphic variants across 92 candidate genes identified by all four approaches of selection screening (Supplementary Table 7), we identified the variant exhibiting the greatest allele frequency difference (*ΔdAF*_cold-warm_) between cold- and warm-region wild boar populations for both regulatory and exonic variants.

For regulatory variants, we identified the highest *ΔdAF*_cold-warm_ in an intronic variant of *IGF1R* (NC_010443.5:g.137677482C>T, c.94+12830G>A, intron 1, rs341219502), demonstrating the highest differentiation of allele frequency (*ΔdAF_cold-warm_* = 0.896). We validated the selection on this variant with the method of extended haplotype homozygosity (EHH) (32, 36). We observed a decay of haplotype homozygosity with increasing distance from the focal core allele (Figure 3a). The EHH decays were far more rapid for the ancestral variant haplotypes (blue curve) than for derived variant haplotypes (red curve, Figure 3b). This signal of positive selection was also supported by the inter-population changes of nucleotide diversity (π) and the level of polymorphism relative to divergence (Supplementary Figure 3a). The nucleotide diversity (π) in the vicinity of rs341219502 was significantly lower in cold-region wild boar populations compared to their warm-region counterparts (the chi-square test, *p* < 1.2 × 10^−4^). Consistently, a significant deficiency of derived polymorphic variants surrounding rs341219502 in cold-region wild boar was found compared to polarized divergent sites at the interspecific level (chi-square test, *p* < 2.2 × 10^−16^). Allele frequency distribution indicated that this variant was absent in tropical Asian populations but was fixed (100%) in northern Asian wild boar populations (Figure 3c). The cross-species orthologous alignment of Amniota vertebrates (Ensembl v105) revealed that the ancestral state ‘C’ is very conserved in *Suidae* species, from *Catagonus wagneri* to 13 pig breeds, indicating that the ‘C’ allele is highly conserved and likely represents the ancestral state (Supplementary Figure 3b). All these findings supported the recent selective sweep in the region surrounding rs341219502 of *IGF1R*.

**Figure 3.**
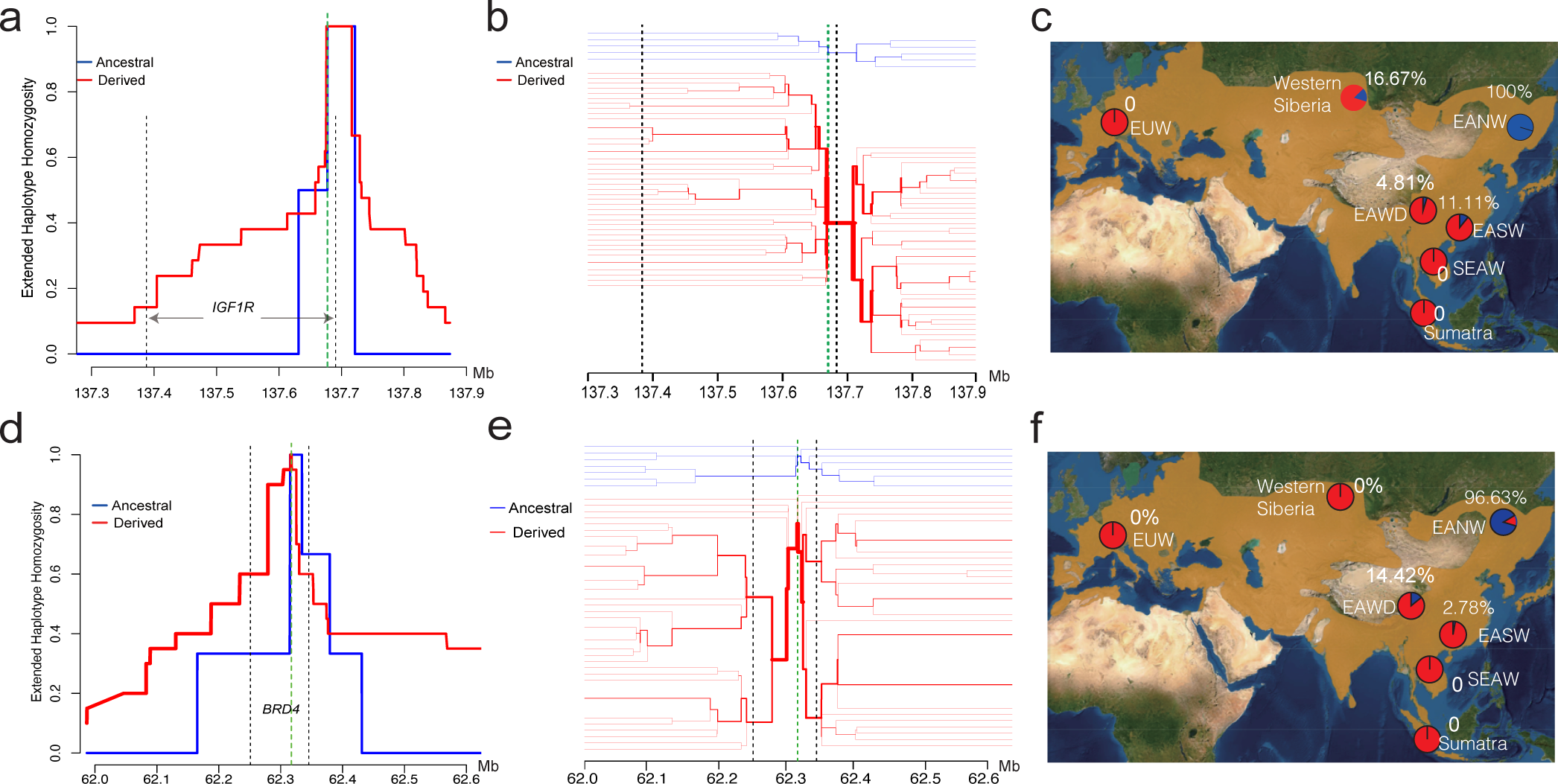
(a) The EHH bifurcation diagram for haplotype density and breakdown around the site rs341219502. Ancestral haplotypes are blue and derived ones are red. The line thickness is positively correlated to the density of haplotypes. (b) The EHH “hat” diagram for ancestral and derived haplotypes around *IGF1R* allele rs341219502. (c) The allele frequency distribution for rs341219502 in multiple populations. (d) The EHH bifurcation diagram for haplotype density and breakdown around the site rs327139795. Ancestral haplotypes are blue and derived ones are red. The line thickness is positively correlated to the density of haplotypes. (e) The EHH “hat” diagram for ancestral and derived haplotypes around *BRD4* allele rs327139795. (f) The allele frequency distribution for rs327139795 in multiple populations.

For exonic variants, we annotated variants with disruptive or protein-altering effects. Based on *ΔdAF*_cold-warm_, we revealed a missense derived variant of *BRD4* (NC_010444.4:g.62317232G>A, c.1043G>A, p.Ser348Asn, rs327139795) with the highest cold-warm differentiation (*ΔdAF*_cold-warm_ = 0.854), among all exonic variants. We observed the decreased nucleotide diversity in the cold-region populations relative to the warm-region populations and the reduced polymorphism relative to the divergence in cold-region populations (Supplementary Figure 3c). We also found the delayed decay of derived haplotype homozygosity (Figure 3d-3e). These findings support the hypothesis of recent positive selection acting on this variant. The nucleotide mutation from ‘G’ to ‘A’ resulted in an amino acid change from Ser to Asn in the second exon of *BRD4*. The multiple species alignment revealed that the ancestral state of the variant ‘G’ is very conserved among mammals (Supplementary Figure 3d). We did not observe the derived allele ‘A’ in nine outgroup species, including African warthogs, pygmy hogs, and *Sus* species, which suggests that the derived allele ‘A’ should have originated after the speciation of *Sus* scrofa. We did not detect the derived ‘A’ allele in wild boar populations from Southeast Asia or Europe, suggesting its East Asian origin (Figure 3f). Among the East Asian wild populations, the derived allele of rs327139795 was nearly fixed in cold-region populations (96.30%) but was a rare allele in warm-region populations (2.78%). The homozygotes of derived allele “AA” were widespread in cold-region samples (92.59%, 25/27) but absent in warm-region populations (0%, 0/18).

### The potential *de novo* origin and recent selective sweep of rs341219502 and rs327139795 in cold-region wild boar

Three candidate scenarios could account for the evolutionary origin of the derived alleles rs341219502 and rs327139795: (1) In the first scenario, the two alleles appeared *de novo* in the cold-region population; (2) In the second scenario, the low-frequency standing variant was transferred from warm-region populations to cold-region populations via gene flow; and (3) In the third scenario, the two alleles appeared *de novo* in the domestic pigs (within last 10,000 years), and then were transferred to cold-region populations via gene flow.

To distinguish between the first two scenarios, we applied gene flow analysis on the genomic region around the focal variants. The localized TreeMix analysis on the genomic regions upstream and downstream of the intronic variant rs341219502 in *IGF1R* (137.3 Mb - 137.6 Mb) indicated gene flow from cold-region populations towards warm-region populations (EANW to EASW in Supplementary Figure 4a). Phylogenetic relationships confirmed this direction of gene flow (Supplementary Figure 4b-c). Specifically, in the background topology of chromosome 1, East Asian wild boar populations were divided into the warm clade and cold clade (Supplementary Figure 4b). However, the local haplotype tree around rs341219502 revealed that five warm-clade haplotypes dispersed into the cold-clade (Supplementary Figure 4c), suggesting the replacement of some warm-clade haplotypes by those from the cold-clade, a typical gene flow process. Thus, the presence of the derived ‘T’ allele in two warm-region wild boar samples likely resulted from southward gene flow from cold-region population. For the exonic variant rs327139795, due to the small number of genic variants (only 21), we expanded our analysis to include broader surrounding regions of the focal variant (61.5 - 63 Mb) and estimated the local gene flow events with TreeMix. The result supported the direction of gene flow from cold- to warm-region wild boar (Supplementary Figure 4d). This observation further suggests that the low frequency of allele ‘A’ in rs327139795 in the warm-region population was most likely introduced into the warm-region population via gene flow from cold regions.

To assess the possibility of the third scenario, we examined the allele frequency distribution among 353 domestic pigs (Supplementary Table 10). We did not detect the derived ‘T’ allele of rs341219502 in either European or southern Chinese domestic pig populations, while being rare variants in East Asian northern and western domestic pigs with low frequencies (2.98% and 4.81%, respectively). Only 2.55% (9/353) of northern Chinese domestic samples carried this derived allele in Min, Meishan, and Tibetan breeds. Moreover, a majority of domestic genotypes (8/9) carrying the derived allele were heterozygous, while only a single Tibetan domestic pig was homozygous. In sharp contrast to this distribution, 93.3% (28/30) genotypes of this allele in cold-region wild populations from Japan, Korea, Siberia, and northern China were homozygous. As for rs327139795, allele ‘A’ homozygotes in European and Chinese domestic populations were also rare (1.07% and 6.58%, respectively). Notably, among all domestic pigs, individuals homozygous for the variant were rare, comprising only 0.28% (1/353) of the population for rs341219502.

Therefore, it is less likely that these two derived alleles originated from domestic pigs. The patterns observed in the origin and rise of rs341219502 and rs327139795 are consistent with the most parsimonious interpretation of a *de novo* origin in wild boar. The alternative hypothesis that these variants originate from ancestral polymorphism is also less plausible because no such variants have been found in Southeast Asian and European wild boars, nor in outgroup species within the Suidae family. Given that the divergence time between Northern and Southern Chinese wild boars falls within 25,000 to 50,000 years ago (37, 38), the age of these two variants could be under 50,000 years. Subsequent to their de novo emergence, natural selection likely facilitated their fixation across wild populations in cold regions.

### The transcriptional changes in rs341219502 of *IGF1R* and post-translational changes in rs327139795 of *BRD4*

Fat and diencephalon are among the tissues responsible for an animal’s ability to withstand cold temperatures (39, 40). Thus, we investigated the expression of *IGF1R* in fat and diencephalon tissues of the Min pig, which is a local breed in cold region of northeastern China. We collected tissue samples from individuals carrying the mutant allele of *IGF1R*. The RNA-seq expression analysis showed that *IGF1R* has high levels of expression in fat and diencephalon of the adult Min pig (Figure 4a). Based on public data of H3K4me1 ChIP-seq for adipose and cerebellum, we revealed that the rs341219502 resides within the enhancer region of the first intron of *IGF1R* (Supplementary Figure 5), suggesting the importance of this variant on regulating gene expression.

**Figure 4.**
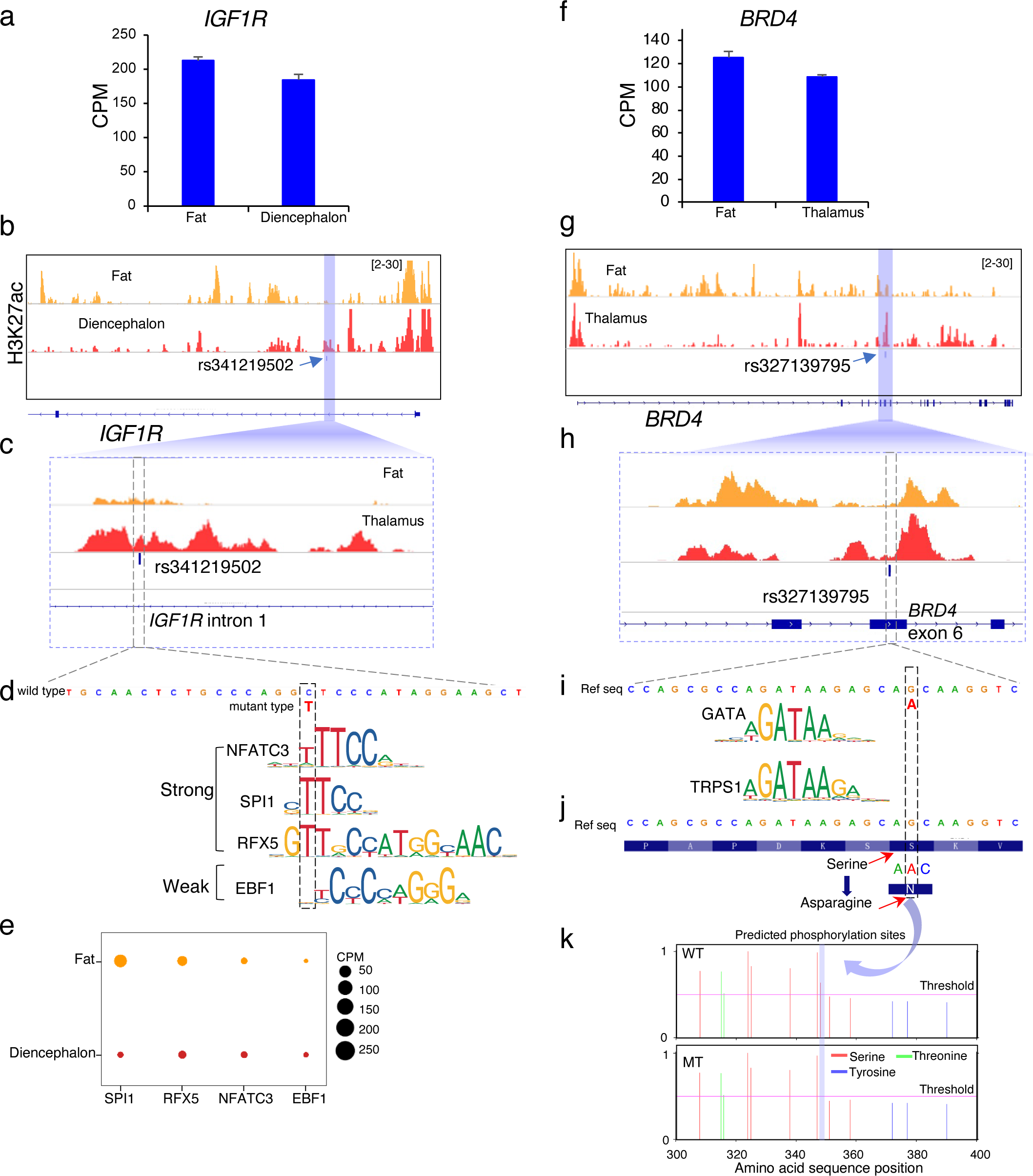
The regulatory enhancer mapping and mutational effect of rs341219502. (a) The RNA-seq expression of *IGF1R* in tissues of fat and diencephalon for the Min pig. (b) The H3K27ac intensity around the gene *IGF1R*. (c) The H3K27ac intensity around the rs341219502 in the *IGF1R* intron. (d) The predicted TF binding sites gained for rs341219502 (C>T) at *IGF1R* intron. (e) The RNA-seq expression profile of TFs in tissues of fat and diencephalon from the Min pig. Note: CPM, or Counts Per Million, is a gene expression normalization to make the expression levels comparable across different samples by accounting for sequencing depth and library size. (f) The expression of *BRD4* in fat and thalamus tissue of Min pig. (g) The H3K27ac intensity around the *BRD4* gene. (h) The H3K27ac intensity around the rs327139795 in *BRD4* exon. (i) The TF binding nearby the rs327139795. (j) The amino acid change of the rs327139795 in exon 6 of *BRD4*. (k) The absence of phosphorylation site in mutant type of rs327139795 in exon 6 of *BRD4*.

Subsequently, we performed the CUT&Tag experiment (41) and found the H3K27ac modification around *IGF1R* (Figure 4b). Specifically, the signals corresponding to the rs341219502 variant were observed within the enhancer region of the first intron of *IGF1R* (Figure 4c). This result confirms the role of this variant on regulating gene expression. The prediction of transcription factor (TF) motif showed that the derived allele ‘T’ could gain three novel TF binding sites, including the NFATC3, SPI1 and RFX5 (Figure 4d). Thus, we analyzed the expression of these three TFs with RNA-seq and found their stable expression in adipose and diencephalon tissues (Figure 4e). These findings indicate that the rs341219502 derived allele ‘T’ is an enhancer mutation, which could potentially enhance the activity of the IGF1R enhancer by introducing novel binding sites for these transcription factors.

We also performed analysis on *BRD4*, a gene also expressed in both adipose tissues and the diencephalon. The H3K27ac modification enrichment were detected around the *BRD4* and its exonic variant rs327139795 (Figure 4f-h). Contrastingly, transcription factor (TF) motif analyses did not identify any TF binding sites at the locus of the *BRD4* exonic variant rs327139795 (Fig. 4i). As this variant resides on the exon of *BRD4*, we further analyzed the amino acid type of allele G and A (rs327139795). The results showed when the genomic sequence changes from G to A, the amino acid at this location (348aa) changes from Serine to Asparagine (Figure 4j). Serine, typically a phosphorylation site, upon substitution to Asparagine, is predicted to result in the loss of this phosphorylation capability. To substantiate this prediction, phosphorylation site analysis of both wild-type *BRD4* and the rs327139795 variant was performed using the NetPhos 3.1 software, which leverages neural network ensembles. The analysis confirmed the loss of the phosphorylation site at the amino acid position 348 upon substitution of Serine with Asparagine (Fig. 5k).

**Figure 5.**
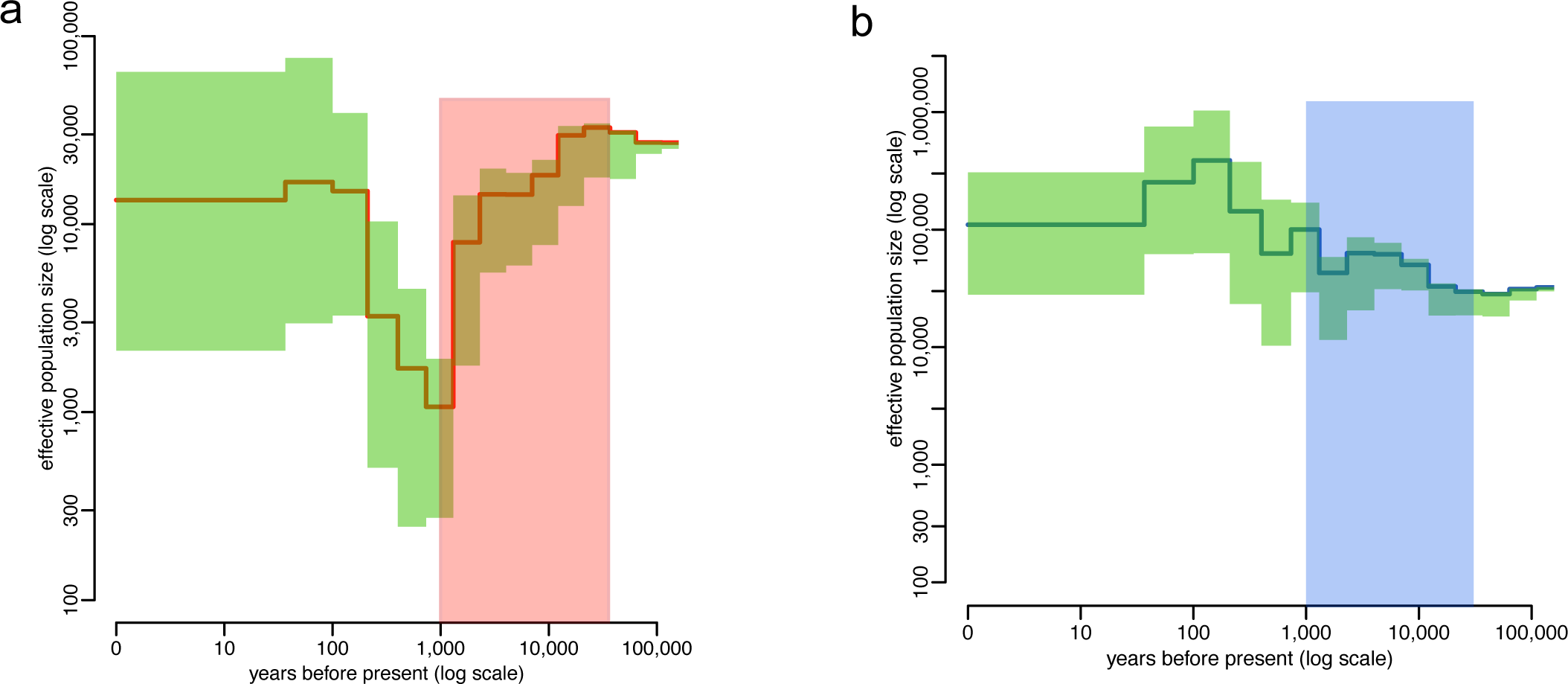
The historical demography of warm-region (a) and cold-region wild boar populations (b) inferred with the approximate Bayesian computation (ABC) based method (30). The dotted lines indicate the 5% and 95% quantiles of the posterior distribution. The red and blue frames show the time range from 50,000 to 1,000 years ago. This period also exhibits the lowest prediction error probabilities (<20%, Supplementary Figure 6). The generation time and mutation rate for simulation were based on a previous study (28).

### The demographic history does not support the ‘genetic drift’ hypothesis

Another question that remains open is: were the selected alleles near-fixed in cold-region populations via the positive Darwinian selection or genetic drift? The fact that we detected strong signals of positive selection on the two sites (rs341219502 and rs327139795) supports the first hypothesis, however, rapid genetic drift can sometimes leave patterns similar to those caused by positive selection (42), so it is necessary to further test the drift hypothesis. As a rapid genetic drift would require a strong reduction in the historical effective population size, we formally evaluated it using PopSizeABC, a simulation-based demographic history inference method operating under the framework of approximate Bayesian computation (ABC). We evaluated the ancestral dynamics of effective population size for both warm- and cold-region wild populations of Asian origin. Based on population-level diploid genomes for warm- and cold-region populations (15 and 21, respectively), 100 independent 2 Mb-long regions, and 600,000 simulated datasets of the same size, we revealed different trends of demographic changes between warm- and cold-region populations ranging from 100,000 years ago to 1,000 years ago. Multiple studies found that the divergence time between Northern and Southern Chinese wild boar is within the range of 25,000 to 50,000 years ago (37, 38), which suggests that wild boar may have arrived in northern China before this period, so this time range covered the upper and lower limits of the divergence time between northern and southern Chinese wild boar. The prediction errors of PopSizeABC inference were within the acceptable range (43) (Supplementary Figure 6). We found a steadily increasing, rather than declining, trend of historical population size for the cold-region wild boar populations during the period of ∼25,000 to 50,000 years ago (Figure 5). In contrast, the population sizes in warm-region wild boar populations decreased sharply during this period. Thus, the increasing trend of population size history in cold-region wild boar does not support the rapid genetic drift hypothesis.

## Discussion

A key challenge in biology is to map out the functional elements within the genome and to understand their roles in adaptive processes. Natural selection can create distinctive patterns in the genomic regions surrounding the positively selected genes, which deviate from those predicted under neutral evolution. Several key indicators— including characteristic variations in genetic diversity, shifts in allele frequencies, and divergences between species—serve as critical tools in the identification of genes under positive selection that are integral to complex adaptations (14, 16, 44-47). For example, research focusing on the plateau wild boar has uncovered genes with selective signals that are vital for coping with severe environmental conditions (47-50). Moreover, the haplotype approach has revealed a significant section of the X chromosome that is involved in climate adaptation in both wild and domestic pig populations from northern China and Europe (16). These studies have substantially deepened our understanding of the dynamics of natural selection.

Northern Asia, encompassing the vast expanse of Siberia, is characterized by its extremely cold winters, posing significant challenges to endothermic mammals in maintaining thermal balance. The severe low temperatures serve as a strong selective force, especially for species like the wild boar, necessitating adaptations for effective thermoregulation in these frigid zones. Beyond the cold, another major environmental challenge is the scarcity of food resources during the lengthy winter season (20). Thus, uncovering molecular adaptations enabling peripheral or derived populations of wild boar to survive and flourish in the cold Siberian climate and its neighboring regions is a subject of great scientific interest.

In this study, employing whole-genome sequencing and four selective sweep scan methodologies, we pinpointed candidate genes that are implicated in cold adaptation. Our pathway analysis indicated several key biological processes, notably in regulating fat cell differentiation, fostering adipose tissue development, thermogenesis, and cold-induced thermogenesis. These insights confirm that brown adipose tissue is essential for thermogenic adaptation to cold, triggering diverse processes involving neural, vascular, and metabolic responses to cold, as revealed from studies in humans, mice, insects, and polar mammals (51-54). Three positively selected genes *(SLCO1C1, PDE3A*, *and TTC28*) were also reported in indigenous Siberian human populations (3). Interestingly, the association between low temperature and SNPs near *SLCO1C1 and PDE3A* has been also found in Holstein cattle (55). Thus, it is likely that these genes are under recurrent convergent evolution in mammals due to their interactive functions for the cold resistance (4, 56).

At gene level, *IGF1R* is particularly interesting considering extensive studies on its gene functions in thermoregulation. Transgenic mice studies have revealed that *IGF1R* can reduce core body temperature under the joint effect of cold stress and calorie restriction (30, 31). *IGF1R* may also have a role in body size regulation (57-59). Among all variants, an intronic variant with most probably regulatory impact (rs341219502, c.94+12830G>A, reverse strand) in *IGF1R* and a missense variant (rs327139795, c.1043G>A, forward strand) in *BRD4* showed the strongest differentiation between the cold- and warm-region populations. Notably, *BRD4* was also among the genes supported by all four methods of selective sweep scans. These two mutations, ‘T’ of rs341219502 and ‘A’ of rs327139795, are absent from all included outgroup Suidae and Tayassuidae species. At the population scale, the alleles are fixed only in cold-region wild boar while rare or absent in warm-region wild boar that are closer to the origin place of pig species (Southeast Asia), which strongly suggests their *de novo* origin in cold-region wild population. Although the two alleles were found in a tiny fraction of the southern Chinese wild boar population and Tibetan domestic pigs, it is most probable that the allele was introduced from northern populations by gene flow, based on the evidence of local TreeMix signals (or phylogeny misplacement), very low allele frequencies, and extremely high heterozygosity rate of allele-carrying individuals from these groups.

Moreover, our CUT&Tag experiment revealed that the derived allele of rs341219502 is located in the enhancer region of *IGF1R*. We validated the enhancer signals of this variant in both fat and diencephalon tissues of the Min pig, a local breed adapted to cold environment of northeastern China. The allele transition from ‘C’ to ‘T’ could result in novel binding targets of TFs, including NFATC3, SPI1 and RFX5, to the enhancer. Interestingly, previous studies showed that NFATC3 is required for cardiac development and mitochondrial function (60). NFATC3 can also increase insulin sensitivity, controlling gene expression influencing the development and adaptation of numerous mammalian cell types including the adipocyte and neurons (61). The SPI1 (PU.1) can inhibit adipocyte differentiation (62, 63). The RFX5 can regulate resistance to nutrient stress (64). Moreover, the ‘A’ allele of rs327139795 within the exon of BRD4 leads to a change from Serine to Asparagine. Based on the basic feature of amino acid, the changing from Serine to Asparagine could affect phosphorylation sites, which may cause changes in the interaction between the mutated Asparagine and surrounding amino acids, thereby potentially affecting protein function.

## Conclusions

The wild boar (*Sus scrofa*) has been remarkably successful in colonizing Eurasia, notably their rapid expansion from tropical Asia into a diverse array of climates. This includes their movement into the extreme cold of arctic Siberia less than a million years ago. Through whole-genome sequencing and employing multiple methods of summary statistics, including the allele frequency spectrum (*F_st_* and the ratio of θ_π_), haplotype, and species divergence, we identified genes under selective sweeps in cold-region populations. These genes were found to enrich metabolic pathways critical for cold resistance, including thermogenesis, fat cell development, and the regulation of adipose tissue. The most pronounced selection signal was identified in a 1.3 Mb region on chromosome 1, characterized by linkage disequilibrium surrounding *IGF1R*. At the variant level, a regulatory variant within *IGF1R* and a missense variant within *BRD4* showed the most significant differentiation in allele frequency between the warm- and cold-region populations. Analysis of allele frequency distribution suggests a de novo origin for these variants within cold-region wild populations. The demographic reconstruction revealed that the rise of these variants was unlikely to be attributable to random effects (such as genetic drift). Considering the known functions of *BRD4* and *IGF1R* in bioenergy homeostasis and body temperature regulation (30, 65, 66), our study provides insight into the molecular adaptation to the cold climate in free-range wild boar populations from Siberia and nearby regions.

## Materials and methods

### DNA sampling, sequencing, variants calling, and datasets

The genomic DNA was extracted from hair follicles of five Vietnamese wild boar (Son La province, Vietnam, ∼20°N), three wild boar from the Novosibirsk region (Novomyhaylovka village, Kochenyovskiy district, Latitude 55°17′35″N, Longitude 81°48′38″E), one wild boar from Tyva (∼51°N), one wild boar from Buryat (∼51°N), and one wild boar from Zabaykalsky Krai (∼52°N). Whole-genome sequencing (WGS) was performed on all samples using the DNBSEQ-T7 platform (MGI) with a paired-end library (2 × 125 bp).

Whole-genome mapping and calling processes were largely conducted following a previously devised methodology (16, 67, 68). In short, whole-genome short-reads data for 26 samples representing wild boar from cold (Siberia and northern Asia, 11 samples) and warm regions (temperate and tropical Asia, 11 samples), and outgroup species (4 samples), were cleaned using fastp software with default parameters (69) and subsequently mapped to the genomic reference of Sscrofa 11.1 with BWA v0.7.17 (70). Although there are multiple reference-level assembled genomes for pigs, including multiple domestic breeds (71, 72) and the wild boar (68), we chose the Sscrofa11.1 as the reference due to its high annotation and sequencing quality (73). For the variant calling, we jointly used two software programs: SAMtools v1.15.1 (74) and GATK v4.2.6.1 (https://gatk.broadinstitute.org/hc/en-us). The major steps included marking duplicates, recalibrating base quality scores, per-sample calling with HaplotypeCaller, and joint-calling with GenotypeGVCFs. We filtered variants using the expression "QUAL < 100.0 || QD < 2.0 || MQ < 40.0 || FS > 200.0 || SOR > 10.0 || MQRankSum < -12.5 || ReadPosRankSum < -8.0".

Two datasets were composed for analyses of specific purposes. The simplified core dataset of 63 samples (Supplementary Table 1) was used for almost all analyses except the initial confirmation of population identity and the allele frequency distribution analysis. For this core dataset, the analyzed samples included 24 wild boar from northern and northeastern China to represent the cold region population, 24 wild boar from southern China, Sumatra, and southern Europe (Italy and Greece) to represent the warm region populations, and 15 samples of four different *Sus* species (*Sus verrucosus, Sus celebensis, Sus cebifrons, Sus barbatus*) as outgroups (16, 75) (Supplementary Table 1). The other dataset was much larger: 488 samples. This dataset was used to confirm the population identity of new samples and evaluate the allele frequency distribution across geographical populations (Supplementary Table 2). This dataset incorporated new samples (11 samples) and the public database data for 477 wild boar and domestic pig samples (Supplementary Table 2). The geographic populations were divided into the European wild boar (EUW, 47 samples), East Asian northern wild boar (EANW, 30 samples), East Asian southern wild boar (EASW, 18 samples), southeastern Asian wild boar (SEAW, 8 samples), and Outgroups species (OG, including African warthog, African bush hog, African red river hog, pygmy hog, Southeast Asian *Sus* species). In addition, we also included domestic pig samples: the European domestic pigs (EUD, 186 samples), East Asian northern domestic pigs (EAND, 84 samples), East Asian southern domestic pigs (EASD, 31 samples), and East Asian western domestic pigs (EAWD, which are also Tibetan pigs, 52 samples).

### Population relationships, ancestry, and gene flow

For the core (63 genomes) and extended (488 genomes) datasets, to understand whether sample size would influence the population relationship, we initially evaluated and compared the phylogenetic topologies of the genomes using a distance matrix derived from identity by state (IBS) calculations. For the core dataset, we also conducted the principal component (PCA) analysis to understand inter-group relationships and population clustering. The potential gene flow between western and eastern Siberian populations was evaluated with the TreeMix analysis (24). We estimated the optimal number of migration events (m) using OptM inference with a sufficient model threshold of 99%(76). We further conducted ancestry estimation and admixture analyses using ADMIXTURE v1.3 (25).

### Genome-wide scan for natural selection signals in the North Asian wild boar populations

We conducted a genome-wide scan for signals of natural selection in North Asian wild boar populations, using four complementary methods: 1. Fixation index (*F_st_*) and nucleotide diversity ratio (θ_π_warm_/θ_π_cold_); 2. The XP-CLR test and nucleotide diversity ratio (θ_π_warm_/θ_π_cold_) (77); 3. Extended haplotype homozygosity (EHH)-based statistic (ihh12) (78, 79); 4. Hudson–Kreitman–Aguadé (HKA)-like test (29, 80).

The fixation index (*F_st_*) and nucleotide diversity (π) are among the most classical parameters, which are based on the expectation that a recent positive selection can reduce the nucleotide diversity but increase the *F_st_* between two populations with differentiated phenotypes. Thus, the selected chromosomal regions for North Asian wild boar are expected to have higher *F_st_* but lower π. The method of ihh12, a haplotype-based scan, was developed to detect both hard and soft selective sweeps, with more power than other tools (such as iHS) to detect soft sweeps (78, 79). The HKA-like test is based on the prediction from the neutral theory that species divergence (fixed site differences) and population polymorphism should be correlated (29). Departures from the strictly neutral evolution via population-level selection would result in a faster reduction of polymorphisms than species divergence (80, 81). We used inter-species differences with numbers of fixed sites (>60 SNPs) as a measure of divergence between *S. scrofa* and other four *Sus* species (*S. verrucosus, S. celebensis, S. cebifrons,* and *S. barbatus*). The XP-CLR test, which is also a cross-population composite likelihood ratio test based on the site frequency spectrum (SFS), is a powerful test for detecting selective events restricted to a single population (77, 82).

The results from these four methods were normalized based on ranked genes and only the genes falling outside of 99% of parameter distribution were considered as significant departures from a strict neutral expectation for each method. Subsequently, to make the identification more rigorous, we considered the genes as positively selected candidates for cold adaptation only if at least three out of four methods supported them independently (Supplementary Table 7). For the positively selected sites, we further confirmed their recent positive selection signal with the method of Extended Haplotype Homozygosity (EHH) around the focal variants (32) implemented in rehh v2 (83). The ancestral-derived relationships were determined based on polarizing sites in the outgroup *Sus* species. The local phylogeny was estimated with FastTree v2.1 (84) based on haplotype consensus sequences composed using SAMtools and BCFtools v1.15 (74).

### Functional pathway enrichment analysis for genes supported by at least three methods and allele frequency distribution for focal variant(s)

Due to the complex nature of climatic adaptation, hundreds of genes would be responsive and under positive selection (3, 4, 56). We thus analyzed functional enrichment to understand the overall pattern of biological pathway. Only genes supported by at least three methods were used for the enrichment analysis with the Metascape database (85). The origin of the derived allele was analyzed based on the allele frequency distribution among populations and across species. The population distribution was based on the extended dataset of 488 samples (Supplementary tables 2 and 10). The focal allele across species was defined as fixed if the homozygous allele was detected in all of the outgroup samples. Ensembl v105 was used to confirm the presence or absence of an orthologous mutation based on the whole genome alignment between *S. scrofa* (Suidae) and a species belonging to the sister family Tayassuidae - *Catagonus wagneri*.

### Historical population demography for cold-region populations

Based on the PCA, ADMIXTURE, and phylogeny, we removed the putative European-origin samples. After filtering out the potential recent gene flow, we kept the closely related samples of Asian origin to represent the cold- and temperate -region populations (15 samples and 21 samples, respectively). In this way, we reduced the impact of current population structure fluctuations on the inference of ancestral dynamics of the effective population size. The computation was based on the instructions of the PopSizeABC pipeline (43). This method applies the framework of approximate Bayesian computation, it can analyze more samples than similar PSMC (86) and MSMC (87) tools, and it is robust to sequencing errors and complex population dynamics (43). Specifically, we first summarized the folded allele frequency spectrum and the average zygotic linkage disequilibrium. Then, we simulated 400,000 datasets under random population size histories. These pseudo-observed datasets (PODs) cover the same sample size and 100 independent 2 Mb-long regions. Ancestral population sizes were independently estimated by the ABC for each POD. The estimated values were compared to their true values with the tolerance rate of 0.001.

### The CUT&Tag and RNA-seq data collection and processing

The samples of Cleavage Under Targets and Tagmentation (CUT&Tag) and RNA-seq were collected from the 180-days Min pig (a Chinese local breed) snap-frozen in liquid nitrogen. All the sample collected protocols were approved by the Ethics Committee of Huazhong Agricultural University (2022-0031). The CUT&Tag experiment followed the guideline of the original CUT&Tag method for tissues (41). The H3K27ac histone modification antibody was used to perform the antibody enrichment of CUT&Tag of fat and diencephalon tissues of the Min pig. The RNA-seq protocols of fat and diencephalon tissues were based on the methods of rRNA-depletion and strand-specific RNA-seq provided by Illumina (Illumina, San Diego, CA). The sequencing for CUT&Tag and RNA-seq was performed with the Illumina NovaSeq6000 (PE150) platform.

The sequencing data of CUT&Tag were cleaned with the Cutadapt to trim the adapter (88). Then the clean data were mapped with Bowtie2 to the susScr11 (Sus scrofa 11.1) reference genome (89). The peak calling was performed using MACS (90). The vertebrate motif from JASPAR2020 were used to match the enhancer sequence of focal genes (91). The RNA-seq data were processed following our previous study (92).

## Acknowledgements

We extend special thanks to Martin Holzenberger and Bruno Conti for their patient discussion and valuable improvements to the manuscript. We also thank Evgeniy Varisovich Kamaldinov, Valeriy Lavrentyevich Petukhov, and our reviewers for their insightful comments and discussions.

## Author Contributions

J.H.C., Y.X.Z., and S.H.Z. supervised this work. J.H.C., M.S., Z.X.X, Y.P.G, R.Z.K, J.Z., Y.Y.J., T.N.T.T., T.S., H.Y, H.M, N.S., J.L.H, D.L, S.Q.X., and I.J. designed the research. J.H.C., J.Z., Z.X.X, Y.P.G, R.Z.K, J.Z., and Y.X.Z analyzed data. J.H.C. wrote the manuscript. Y.X.Z, S.H.Z, and I.J. revised the draft.

## Data Availability Statement

The whole-genome sequence data can be accessed through NCBI BioProject code PRJNA859556.

---

**Supplementary Figure 1.** Gene flow analyses among major populations (a) The cross-validation errors of the ADMIXTURE tool for inferring population ancestry and admixture. (b) The optimal number of migration events (m) for TreeMix based on the inference of OptM estimation. Over 99.8% of the variance was explained when m=3. (c) The δm estimation supported the migration events of 3. (d) The direction of gene flow revealed by TreeMix, based on autosomal SNPs. The gene flow from eastern to western Siberia was detected (EFG → H). Three arrows show the directions from donor populations to recipient populations. Note: all the population codes are the same as in Figure 1a.

**Supplementary Figure 2.** The warm-cold nucleotide diversity (π) comparison and HKA test (p<0.01) for the neighboring genes *SLCO1C1* and *PDE3A* (a) and *TTC28* (b). Dotted lines indicate gene boundaries.

**Supplementary Figure 3.** (a) The local signals of *IGF1R* selective sweep based on evidence of nucleotide diversity (π) for cold- and warm-region populations (left axis) and the polarized HKA test (right axis). The red arrow indicates the site of rs341219502. (b) The population and evolutionary conservation for the ancestral state of rs341219502 (“C”) based on the whole-genome alignment. The first sequence represents the mutant sequence of cold-region wild boars. The next 13 sequences are retrieved from genomes of pig breeds in Ensembl (v105) and the last sequence shows *Catagonus wagneri* genome (Ensembl v105). (c) The local signals of *BRD4* based on evidence of nucleotide diversity (π) for cold- and warm-region populations (left axis) and the polarized HKA test (right axis). The red arrow indicates the site of rs327139795. (b) The population and evolutionary conservation for the ancestral state of rs327139795 based on the whole-genome alignment.

**Supplementary Figure 4.** The derived allele frequency distribution and the gene flow direction. (a) The localized gene flow around variant rs341219502 (600 Kb) revealed by TreeMix. Three arrows show the directions from donor populations to recipient populations. All population codes are the same as in Figure 1a. (b) The background topology of chromosome 1 with a highlight on the two clades of North-region (cold) and South-region (warm) populations, represented by blue and red branches, respectively. (c) The local haplotype tree of 600 Kb around the rs341219502 inferred with the Maximum likelihood method of FastTree v2.1. The red haplotypes from the warm-region population were nested into the clade of the cold-region population. The “1” and “2” represent the two haplotypes for each sample. Note: The black asterisks indicate major clade branching with 100% support values. The triangles represent wild boar samples from temperate or tropical regions that are nested within the clade of the cold region. The abbreviations of regions are: EUW, European wild boar; EAWD, East Asian western domestic pigs (Tibetan breed); SEAW, southeastern Asian wild boar; EASW, East Asian southern wild boar; EANW, East Asian northern wild boar (including populations from northern China, Korea, eastern Siberia, and northeastern China). (d) The localized TreeMix migration events (m=3) for the region from 61.5Mb to 63Mb around the 811 selected variant rs327139795 of *BRD4*. The directions of gene flow are shown with arrows.

**Supplementary Figure 5.** The mapping signals based on H3K4me1 and H3K27ac ChIP-seq data retrieved from UCSC Genome Browser (http://genome.ucsc.edu/s/zhypan/susScr11_15_state_14_tissues_new). The red arrow shows the coordinate to the intronic variant rs341219502 of *IGF1R*.

**Supplementary Figure 6.** The predicted errors for different time stages. The time range from 50,000 to 1,000 years ago has the lowest errors (<20%, scaled) under the tolerance rate of 0.001. The red and blue lines show the warm- and cold-region populations, respectively.

