## Supplementary figures and images for "Positive Selection on Rare Variants Underlying the Cold Adaptation of Wild Boar"

### Supplementary Figure 1

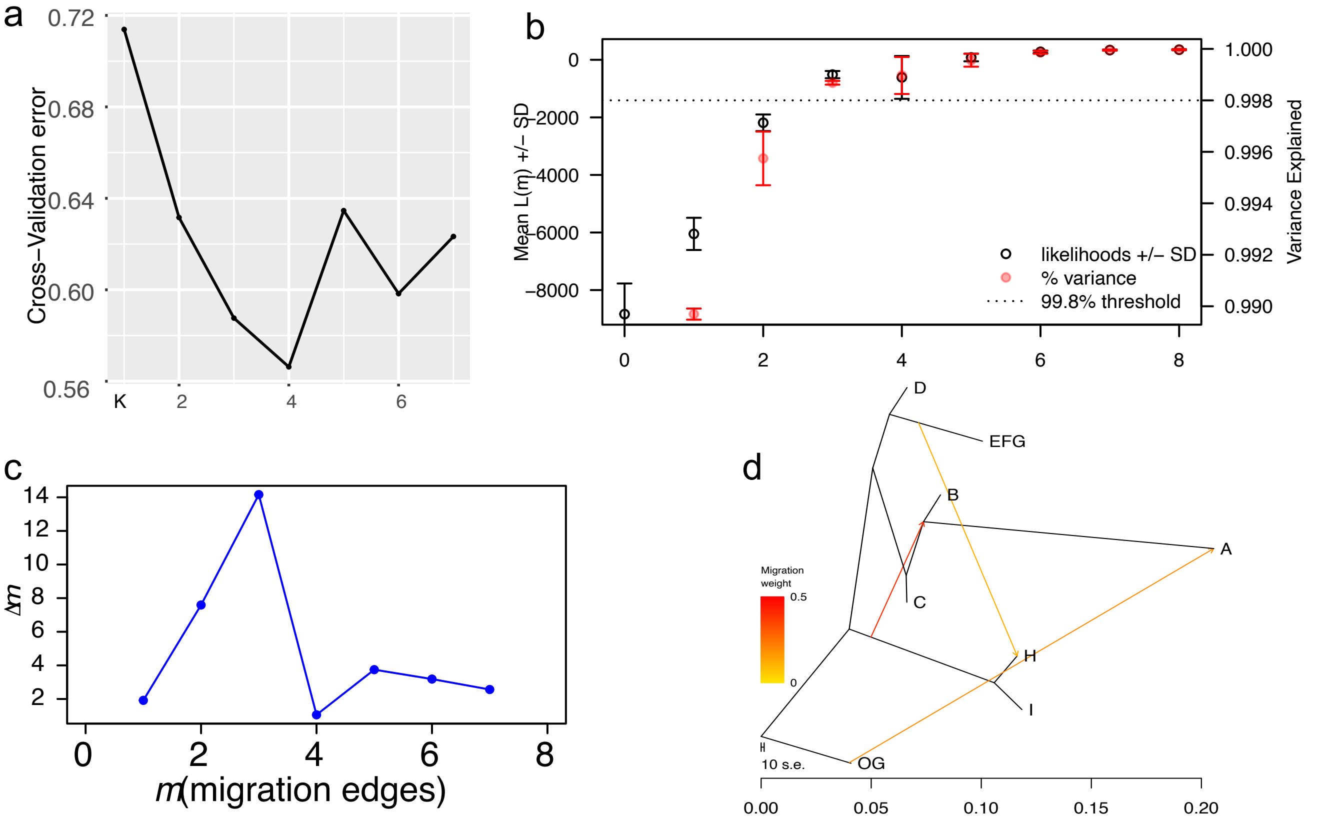

### Supplementary Figure 2

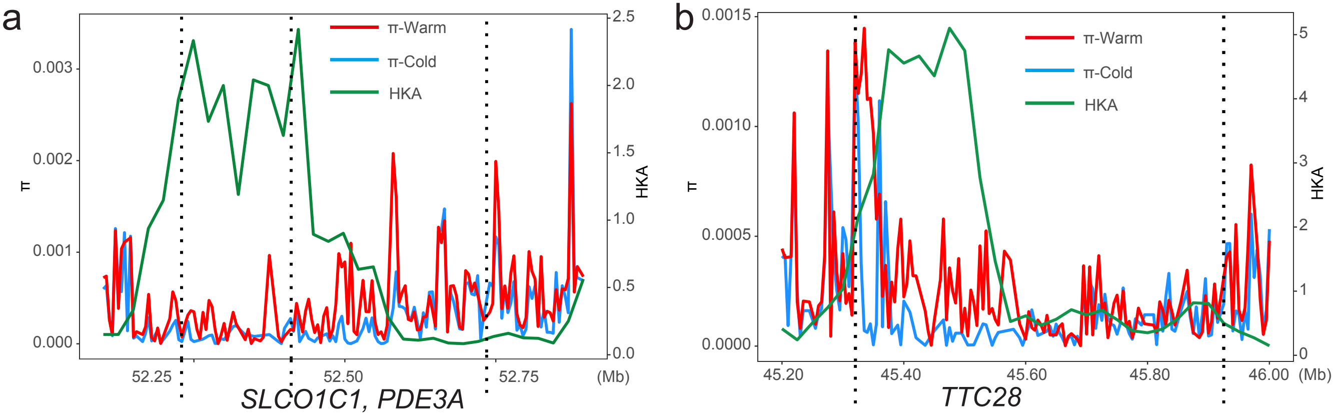

### Supplementary Figure 3

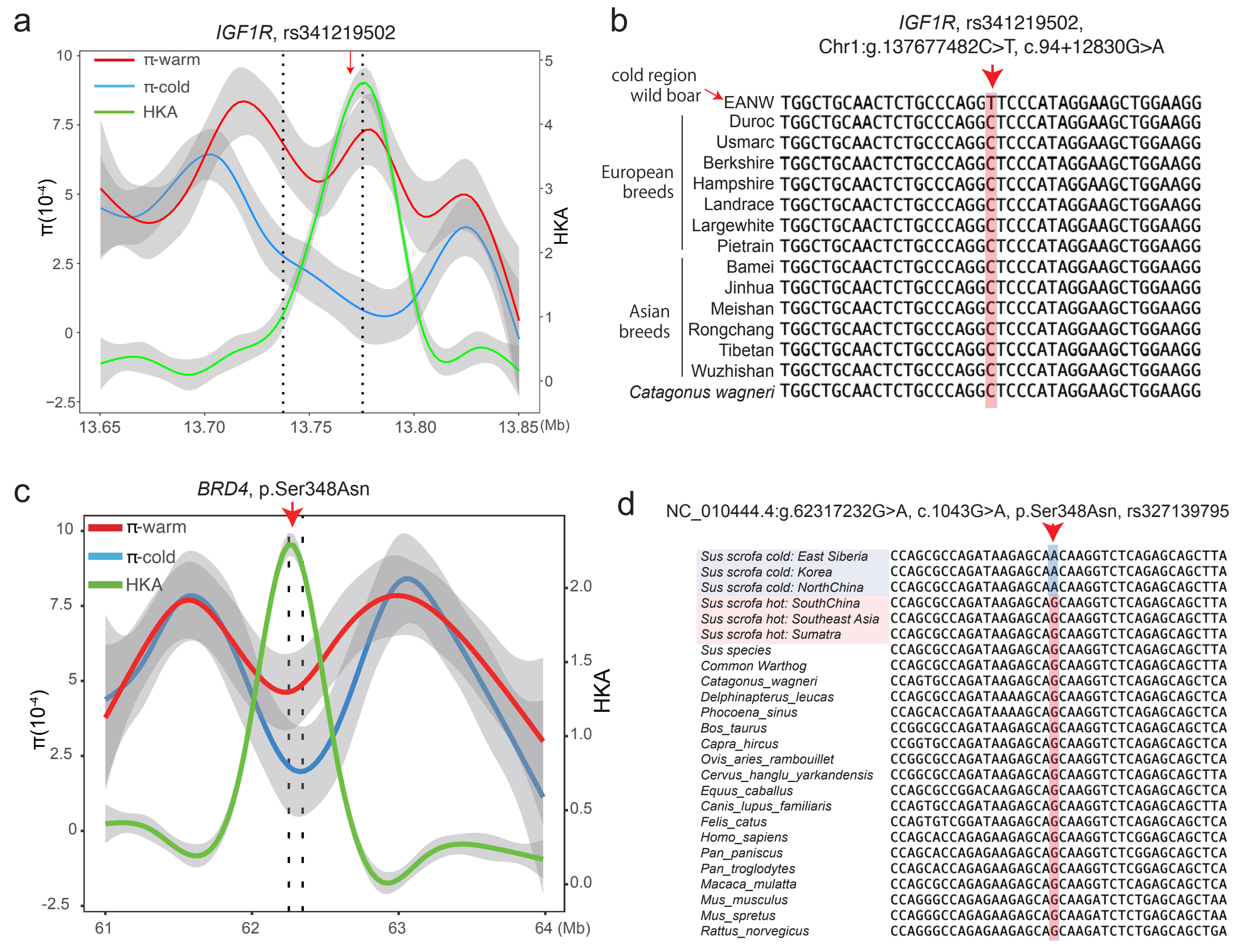

### Supplementary Figure 4

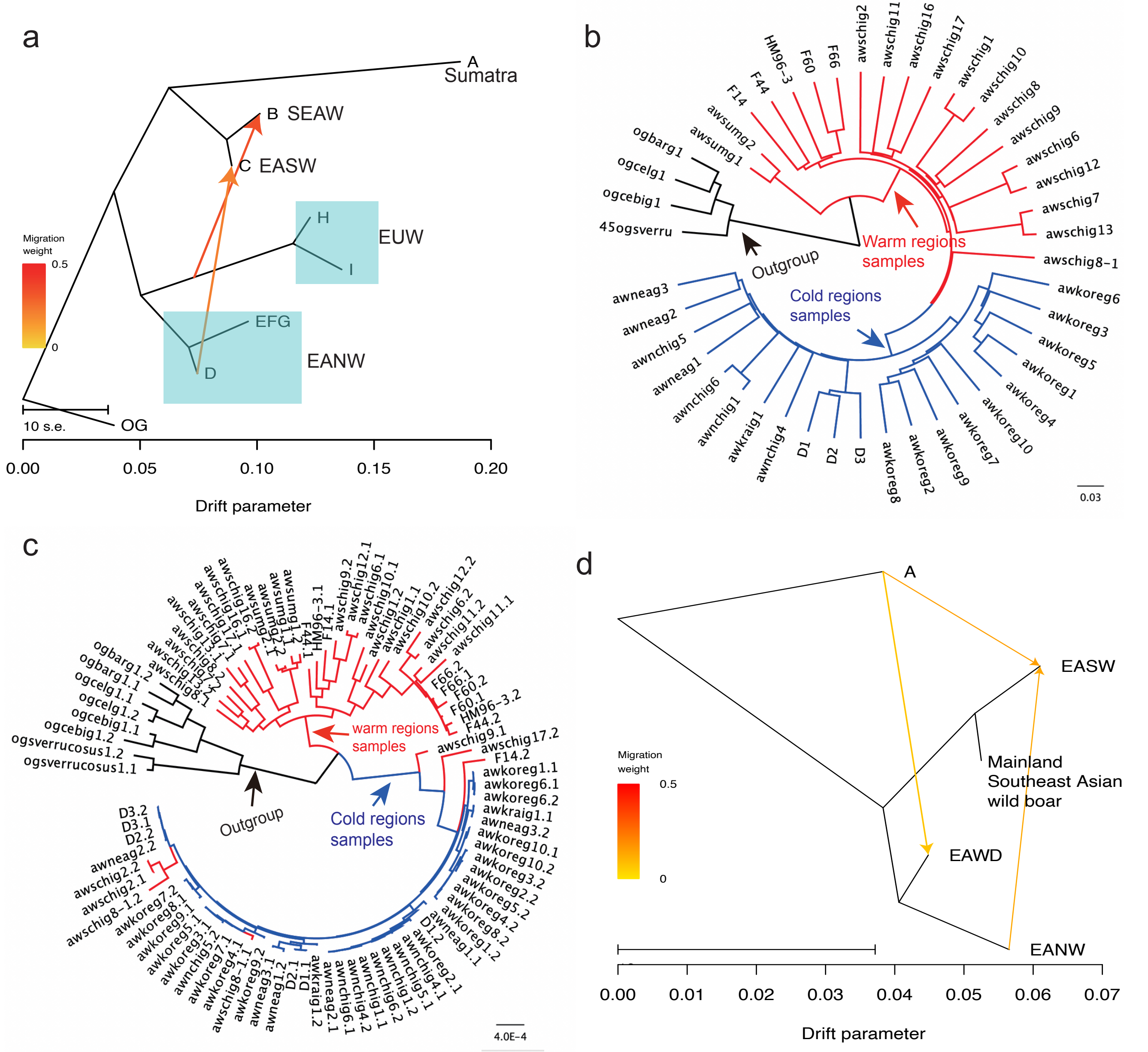

### Supplementary Figure 5

NC_010443.5:g.137677482C>T, c.94+12830G>A, rs341219502

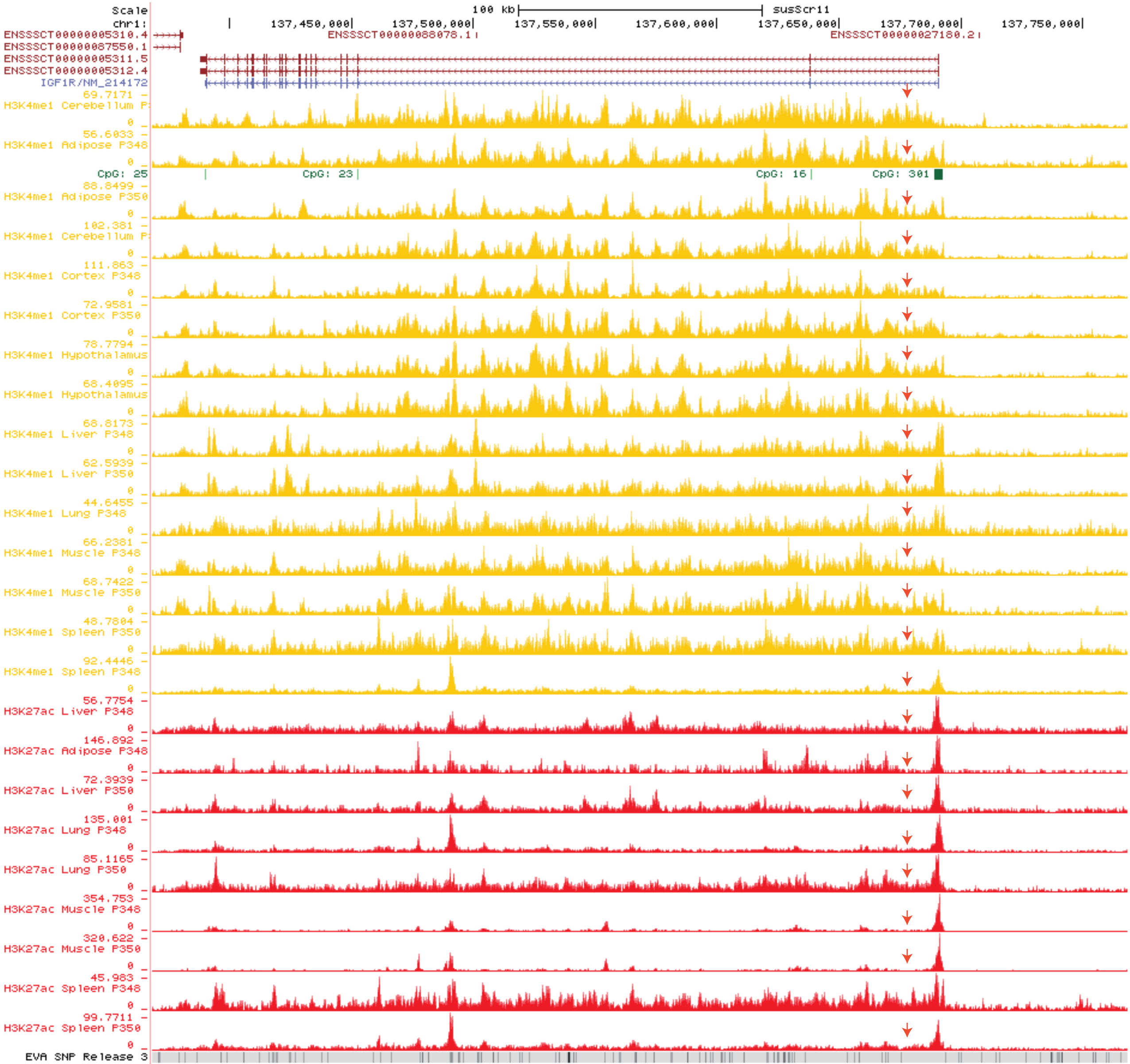

### Supplementary Figure 6

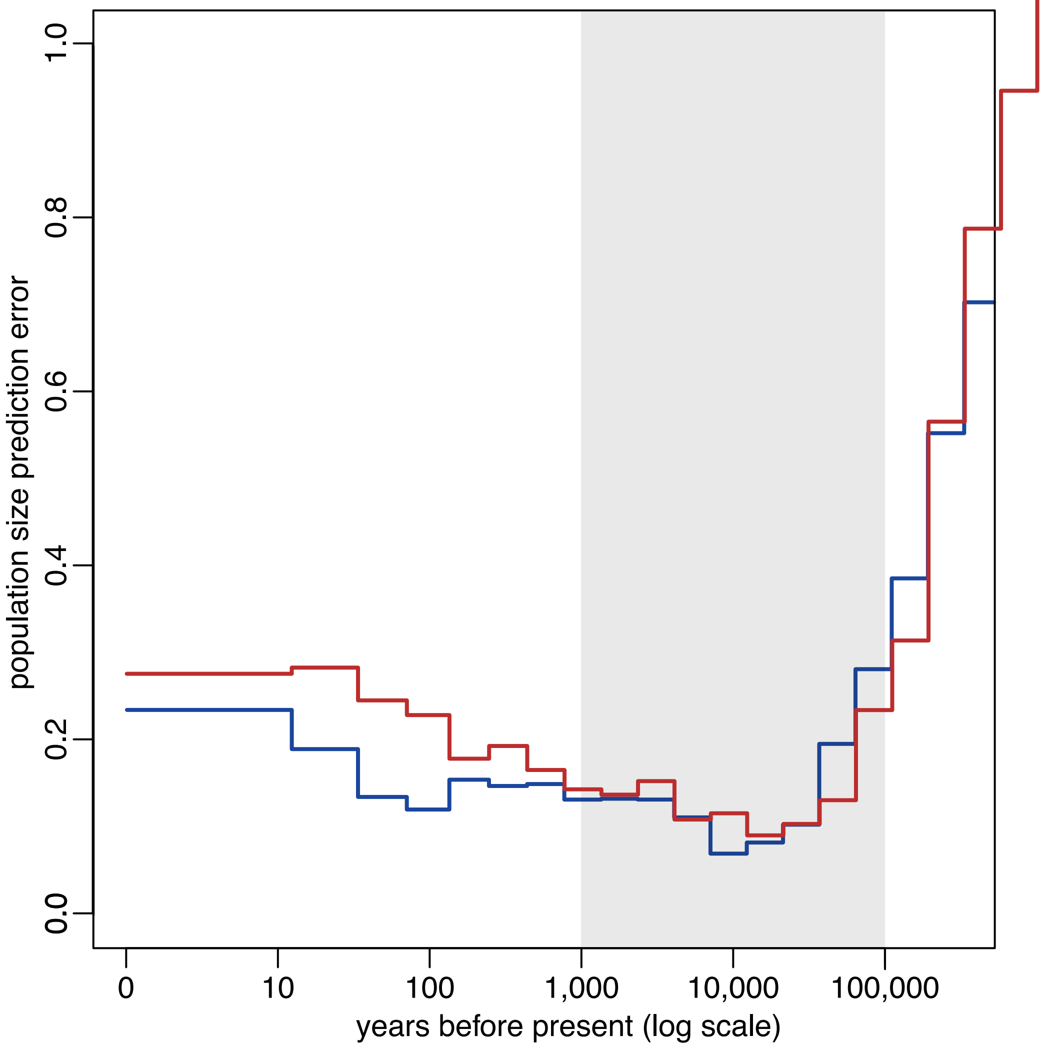
